# Cool temperature acclimation in toxigenic *Microcystis aeruginosa* PCC 7806 and its non-toxigenic mutant

**DOI:** 10.1101/2023.08.28.555099

**Authors:** Gwendolyn F. Stark, Robbie M. Martin, Laura E. Smith, Bofan Wei, Ferdi L. Hellweger, George S. Bullerjahn, R. Michael L. McKay, Gregory L. Boyer, Steven W. Wilhelm

**Affiliations:** Department of Microbiology, The University of Tennessee, Knoxville, TN, USA; Department of Chemistry, State University of New York College of Environmental Science and Forestry, Syracuse, NY, USA; Water Quality Engineering, Technical University of Berlin, Berlin, Germany; Department of Biological Sciences, Bowling Green State University, Bowling Green, OH, USA; Great Lakes Institute for Environmental Research, The University of Windsor, Windsor, ON, Canada

**Keywords:** cyanobacteria, cyanotoxins, transcriptomics, chemostats, microcystin

## Abstract

For *Microcystis aeruginosa* PCC 7806, temperature decreases from 26° C to 19° C double the microcystin quota per cell during growth in continuous culture. Here we tested whether this increase in microcystin provided *M. aeruginosa* PCC 7806 with a fitness advantage during colder-temperature growth by comparing cell concentration, cellular physiology, and the transcriptomics-inferred metabolism to a non-toxigenic mutant strain *M. aeruginosa* PCC 7806 Δ*mcyB*. Photo-physiological data combined with transcriptomic data revealed metabolic changes in the mutant strain during growth at 19° C, which included increased electron sinks and non-photochemical quenching. Increased gene expression was observed for a glutathione-dependent peroxiredoxin during cold treatment, suggesting compensatory mechanisms to defend against reactive oxygen species are employed in the absence of microcystin in the mutant. Our observations highlight the potential selective advantages of a longer-term defensive strategy in management of oxidative stress (*i.e.,* making microcystin) *vs* the shorter-term proactive strategy of producing cellular components to actively dissipate or degrade oxidative stress agents.

**Importance:** Through comparisons of a microcystin-producing wildtype strain *M. aeruginosa* PCC 7806 and a non microcystin-producing mutant, *M. aeruginosa* PCC 7806 *ΔmcyB*, our observations highlight defensive (microcystin production) *vs* active (production of degradation enzymes and increased electron sinks) strategies for dealing with cold-temperature induced oxidative stress as well as associated physiological changes. This work increases our understanding of microcystin’s intracellular function, and the role it may play in bloom ecology. In combination with other studies, this work begins to experimentally establish a mechanistic foundation to better understand cold-to-warm seasonal transitions from toxigenic to non-toxigenic strains frequently observed *in situ*.

## Introduction

*Microcystis*-dominated harmful algal blooms (HABs) occur throughout the summer season. For HABs that occur in temperate North American climates, such as Lake Erie, there are seasonal trends between temperature conditions experienced in early June, when temperatures can average 18° C, compared to those recorded later in the bloom season (August), when surface water temperatures can reach 26° C (1, 2). As the season progresses and temperatures warm, harmful algal blooms in Lake Erie have shown a trend of becoming less toxic in conjunction with a shift from toxigenic-to non-toxigenic genotypes (1, 3, 4). The drivers of this genotypic change still remain unknown (5).

We previously observed that short-term temperature decreases (within the span of a week) can induce a “cool-temperature phenotype” in *Microcystis aeruginosa*, whereby microcystin production increased (2, 6). Past research on HAB responses to temperature have largely focused on temperature increases due to concerns for a changing global climate (7). However, when considering cold-to-warm seasonal bloom dynamics, understanding microcystin production across seasonal growth temperatures is relevant. Additionally, lakes subject to cooler stream discharge or which receive inputs from glacial or snow water melt can experience reductions in their thermal structure even in warmer months (8). Changes in microcystin production during growth at cooler temperatures remains a critical component in understanding *Microcystis* bloom ecology (2, 6).

While regulation of the cold stress response in cyanobacteria is complex, both the physical properties of the thylakoid membranes (fluid or rigid) and the redox state of the plastoquinone pool (PQ pool) are involved in either sensing or responding to cold stress (9–12). In cyanobacteria, the thylakoid membranes house both the photosynthetic and respiratory electron pathways, which converge at the PQ pool, making its redox status important for regulating electron transfer and energy dissipation processes in the cell (13–17). Microcystins have been implicated to have a redox-related function. Research has shown differences in redox-related protein abundances between toxigenic and mutant strains of *Microcystis aeruginosa* during iron-limitation (18). Furthermore, thiol-groups, important in maintaining redox balance through glutathione and thioredoxin networks, bind to microcystins *in vivo* (19, 20). The redox state of the of the photosynthetic apparatus in *Microcystis* grown under different light intensities was also negatively correlated with the microcystin content per cell (21). Conclusions drawn from these studies suggest microcystin could play a role in redox-mediated metabolism.

Thiol-groups play a role in redox balance, where the presence of a reactive cysteine residue (thiolate) subjects them to post-translational modifications from reactive oxygen (ROS) and nitrogen species (RNS) (22, 23). This property makes thiol-groups important for redox signaling and enzymatic redox regulatory pathways (24). The original hypothesis offered to explain the binding of microcystin to redox-sensitive thiol groups was that it protected proteins and stabilized them from protease degradation/oxidation (19). However, the idea of *in vivo* stabilization was not directly substantiated. While binding of microcystin to thiols could play a stabilizing role, another possibility is that binding of microcystin to proteins may interfere with repair or regulatory functions in the cell (25, 26). Microcystins are known to form a covalent linkage with a key thiol in protein phosphatase 1A in eukaryotes, thereby permanently inactivating these important regulatory proteins (27). Similarly, microcystin has been shown to bind to phycobilisome subunits (19). It was hypothesized this binding might serve to stabilize the phycobilisomes from protease degradation. However, in nutrient-limiting conditions, phycobilisome degradation *via* proteases can be beneficial, therefore microcystin binding might instead interfere in these processes (28, 29). In these situations, binding of microcystin to proteins might alter metabolic activity that is reliant on protease-mediated degradation or on regulation mediated by the reduction/oxidation of thiol-groups. These interactions alone would mean that microcystin could have a multi-functional regulatory role in the cell, which could help explain why various environmental factors can influence microcystin production (30).

In continuous cultures of *M. aeruginosa* PCC 7806, a decrease in temperature from 26° C to 19° C induces a two-fold increase in microcystin quota (6). Mechanistically, we hypothesized this response was linked to excitation pressure and the production of oxidative stress as a result of constant light at colder temperatures (cf. 31). In our previous study, peak microcystin quotas coincided with cell recovery and the establishment of a new steady-state cell concentration at 19° C. It was unclear if increased microcystin quota was a key acclimation mechanism, and to what extent other mechanisms were involved. Here we tested the effects of cool temperature on *M. aeruginosa* PCC 7806 *vs.* the microcystin-free mutant *M. aeruginosa* PCC7806 𝚫*mcyB* (32). Through comparisons of transcriptional and physiological responses, we identified alternative mechanisms employed by the mutant to acclimate to 19° C. Our observations support the idea that in the absence of microcystin as a presumed physiological defense against oxidative stress, cells actively work to dissipate energy and degrade oxidative stress compounds that are generated as byproducts of excess light adsorption during cold stress. In *Microcystis*, the trade-offs between defensive *vs.* active approaches in mediating oxidative stress establish a potential mechanistic foundation, as well as a new hypothesis, for the regularly observed seasonal transition from toxigenic to non-toxigenic strains during bloom succession.

## Results

### Cell concentration in chemostats

At 26° C, wildtype steady-state cell concentration averaged 3.07 x 10^6^ mL^-1^ (Fig. 1A). When temperature was lowered to 19° C, growth rate dropped, causing cell concentration to decrease to an average of 1.76 x 10^6^ mL^-1^. By T5 (168 h), wildtype cell concentration stabilized around 2.04 x 10^6^ mL^-1^ for the remainder of the cold treatment (T6 through T8). In the mutant strain, a similar trend in growth rate decline and cell concentration was observed (Fig. 1A). At 26° C, the mutant steady-state concentration averaged 2.46 x10^6^ mL^-1^. Temperature change to 19° C resulted in an average concentration decline to 1.27 x10^6^ mL^-1^. By T6 (∼192 h), cell concentrations stabilized around 1.39 x 10^6^ mL^-1^ for the remainder of the cold treatment. When temperature returned to 26° C at T9, wildtype cell concentration increased to an average of 2.12 x 10^6^ mL^-1^ over 24 h, while the mutant cell concentration increased to an average of 1.88 x 10^6^ mL^-1^ at T9 (336 h).

**FIG 1.**
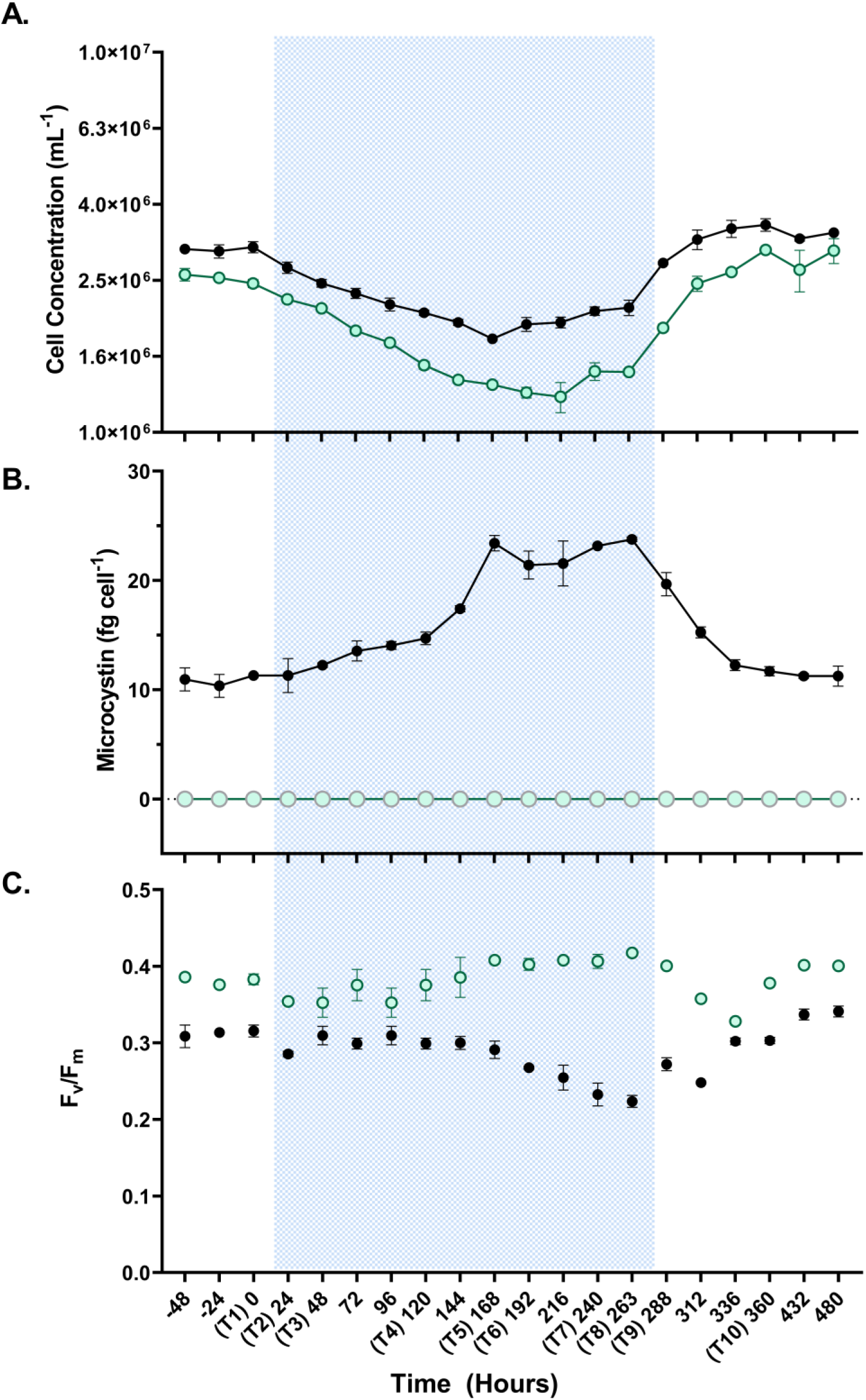
*Microcystis aeruginosa* PCC7806 wild-type and mutant responses to temperature change in chemostats. (A) Average cell concentrations from duplicate wildtype and mutant mono-culture chemostats. Mutant strain is represented by the green open circles, wildtype is represented by black closed circles. Blue shading indicates the 19°C cold period, white is indicative of the 26°C warm period. The x-axis is in units of hours, with each of the ten RNA-seq time points. (B) Average microcystin quota in fg/cell from the duplicate wildtype and mutant monoculture chemostats. The mutant strain had no detectable microcystins. The wildtype microcystin quota roughly doubled after 168h in cold treatment, which corresponds to RNA seq time point T5. The increased microcystin quota persisted in the wildtype for the rest of the cold treatments (T5-T8). (C) Average maximum quantum yield of PSII (F_v_/F_m_) from duplicate wildtype and mutant monoculture chemostats. The mutant shows a steady significant increase in F_v_/F_m_ during cold treatment by T8 in comparison to T1 (p= 0.0461). The wildtype strain F_v_/F_m_ shows an inverse trend from the mutant, significantly decreasing by T8 compared to T1 (p < 0.0001). In response to the temperature change back to 26°C, the mutant F_v_/F_m_ significantly declined at 312h (p=0.044) and 336h (p=0.004), compared to F_v_/F_m_ taken at T9.

### Cellular microcystin content

Microcystin concentrations in wild-type cells were highly reproducible between the paired chemostats (largest difference ± 17%) and reached steady-state concentrations of ∼10 fg cell^-1^ at 26° C (Fig. 1B). After temperature decrease to 19°C, toxin concentration peaked at T5, reaching 23.4 fg cell^-1^. Microcystin remained elevated (>20 fg cell^-1^) in the wildtype for the duration of the cold period (T6-T8). When temperature returned to 26° C, toxin declined to pre-cold-treatment levels. No microcystins were detected in *ΔmcyB* cultures (Fig. 1B). Though no toxin was produced in the mutant, transcription of *mcyD* and *mcyA* followed the expression pattern of the wildtype strain throughout our RNA-seq time series (Fig. S1A-B).

### Photosynthetic physiology assessment

In the wildtype, F_v_/F_m_ values stayed consistent throughout the cold treatment, until T6 (∼192 h) (Fig. 1C). At T6, F_v_/F_m_ values decreased, becoming significantly lower at T7 (∼240 h) (p = 0.038) and T8 (∼263 h) (p < 0.0001) compared to T1 (0 h). In the mutant, F_v_/F_m_ was consistently higher than the wildtype, regardless of treatment (Fig. 1C). During the cold treatment, F_v_/F_m_ in the mutant was higher at T6 (p= 0.016) and T8 (p= 0.046) than at T1 (Fig. 1C).

The mutant and wildtype responded differently during re-acclimation to 26° C. F_v_/F_m_ declined significantly in the mutant in the 48-h period after the temperature was returned to 26° C (T9 vs. 312 h, p = 0.044; T9 vs. 336 h, p = 0.004) (Fig. 1C). In contrast, in the wildtype, F_v_/F_m_ increased in the same 48-h period, but the increase was not significant compared to T9.

Non-photochemical quenching (NPQ) measurements were derived from the rapid light curves (Fig. 2). In the wildtype, there were no significant differences in NPQ measurements during the cold treatment (T5) compared the control (T1) (Fig. 2). In the mutant, there was a significant increase in NPQ measurements taken during the cold treatment (T5) compared to the control (T1). These increases occurred at all PAR (μmol photon m^-2^ s-1) values >345 (Holm-Sidak p_adj_ all <0.05) (Fig. 2, Fig. S2). At T1, the mutant had higher NPQ than the WT across all PAR intensities measured, but the differences were not significant. However, NPQ was significantly higher in the mutant at T5 at all seven measured PAR values (p= 0.024, 0.005, 0.001, 0.006, 0.008, 0.005, 0.003). These data suggest differences in acclimation strategies between the two strains.

**FIG 2.**
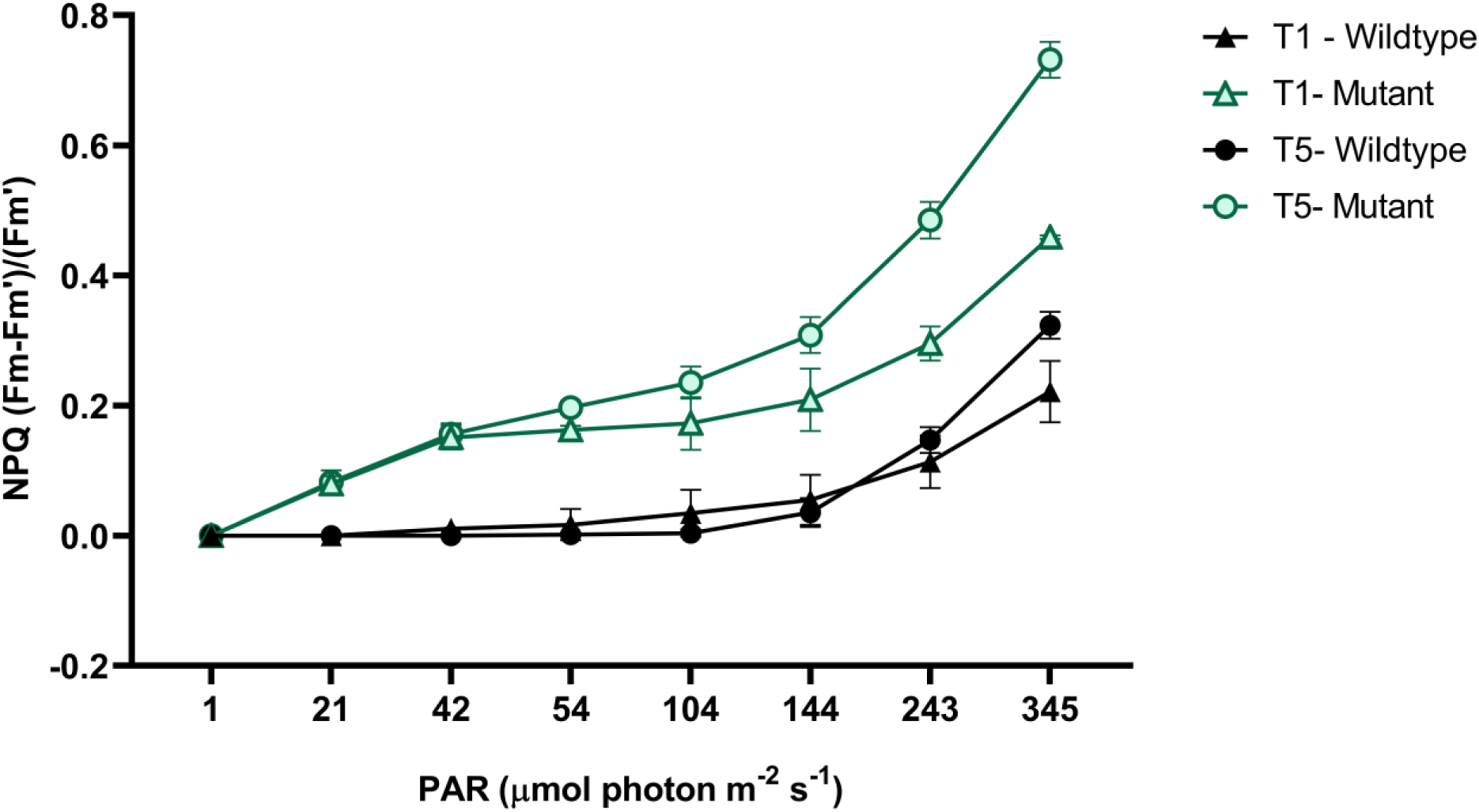
Average NPQ collected from the RLC for each chemostat, at T1 (0 h) and T5 (∼168 h). NPQ significantly differs between the mutant and wildtype for measurements taken at the T5 time point across all actinic PAR measured (Holm sidak adj. (p-values = 0.024, 0.005, 0.001, 0.006, 0.008, 0.005, 0.003).

To assess stress on photosystem II at T5 versus control timepoints (T1 and T10), excitation pressure measurements were derived from the rapid light curves (Fig. S3). The wildtype had significantly higher excitation pressure at T5, compared to T1 and T10 (p= 0.0161, 0.0120). The mutant had significantly higher excitation pressure at T5 versus T10 (p= 0.0087).

### Reactive oxygen species damage measurements via the CM-H_2_DCFDA Assay

The CM-H_2_DCFDA assay revealed significantly higher net green fluorescence (p < 0.0001) in the mutant compared to the wildtype strain during the eleven-day cold temperature treatment period (Fig. S4A-B). There was no significant difference in net green fluorescence between the strains during growth at 26°C.

### Transcriptional profiles for Control, Cold Treatment, and Cold-Recovery

The transcriptional profiles of chemostat populations from ten time points is provided in Table S1. We sequenced 40 mRNA libraries with total reads ranging from 25,573,750 to 78,079,140 per library (Table S2). Broad transcriptional patterns were determined by projecting samples in two-dimensional space (nMDS) based on the expression of all genes in each genome (Fig. 3). This nMDS representation resulted in a circular progression across measured time points (T1-T10). In both strains, T2-T9 scale away similarly from T1 in a counter-clockwise pattern, showing an increasing divergence in gene expression during cold acclimation. However, the mutant transcriptomes diverge from T1 more than the wildtype. Re-acclimation to 26° C (T10) resulted in the transcriptional profile returning to a state similar to T1 in both strains.

**FIG 3.**
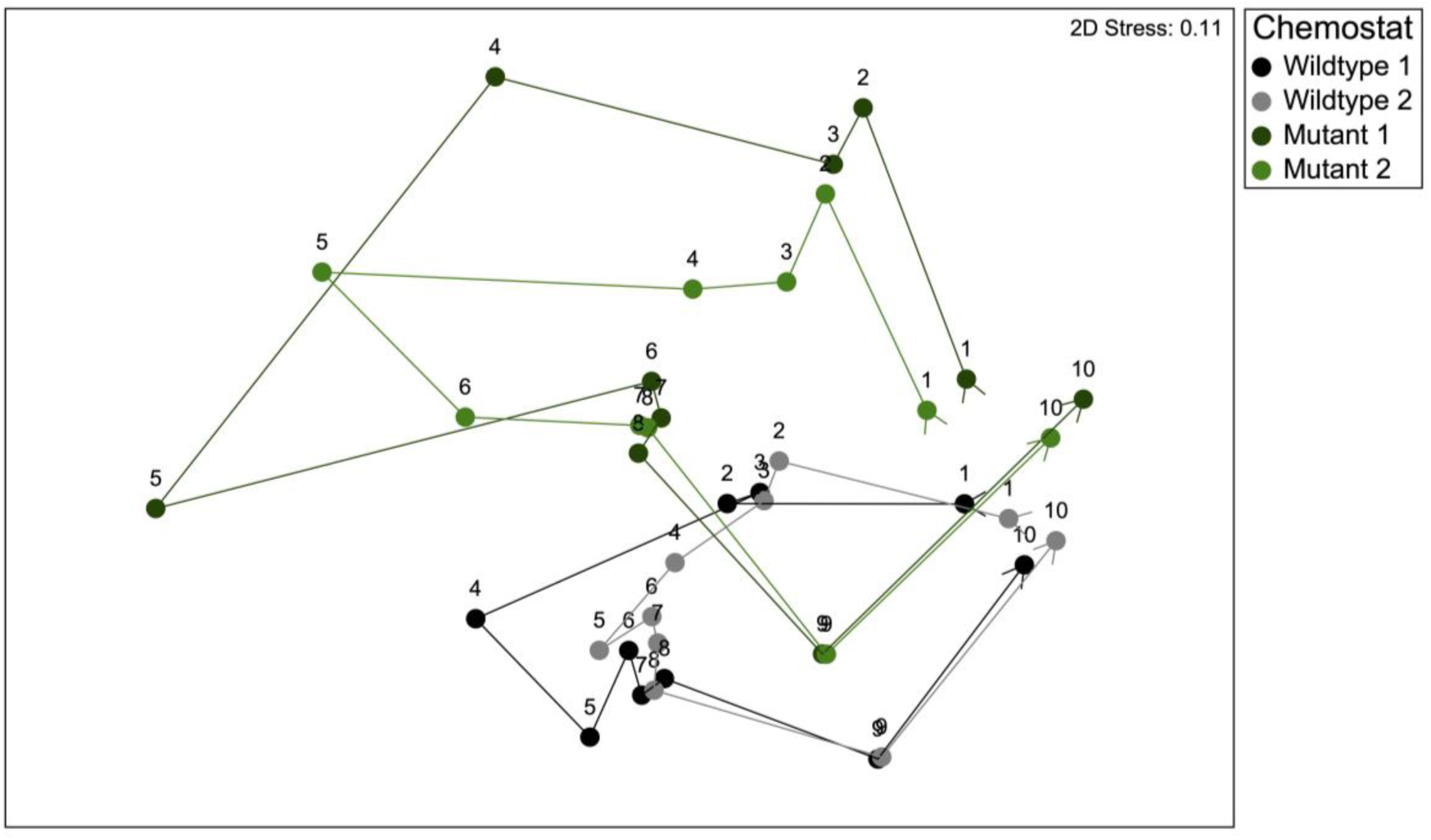
NMDS of square root transformed TPM data encompassing the whole transcriptome for wildtype (black and gray circles) and mutant (green and dark green circles) monoculture chemostats, n=40. The nMDS was constructed to see how the transcriptomes changed over time throughout the cold temperature treatment at each RNA-seq timepoint (T1 through T10).

*Transcriptomic differences between the mutant and wildtype in warm conditions (T1 and T10).* There were 61 differentially expressed genes (fold change ≥2) at T1 and 94 at T10, many of which (42 genes) overlapped. A significant finding was differential expression of genes located in a putative gene cluster (gene cluster 1) with homology to transmembrane proteins, and genes with functions related to redox metabolism/membrane potential (Table S13). At T1, all genes from gene cluster 1 were upregulated 4.6 to 22.8-fold (all p < 0.001) in the mutant strain (Fig. S5, Table S3). At T10, transcripts from seven of the genes in putative gene cluster 1 were upregulated 2.3 to 9.1-fold in the mutant (p < 0.001) (Table S12). All other differentially expressed genes at these time points (T1 and T10), and the rest of the RNA seq time points (T2-T9) can be found in Tables S3-S12.

### Transcript differences between the mutant and wildtype in cold conditions (T2-T8)

Generally, differences in gene expression during cold growth involved changes in a glutathione-dependent peroxiredoxin, and cytochrome-based electron transport pathways (Fig. 4, Fig. 5A-D). Cold treatment produced differential expression of a glutathione-dependent peroxiredoxin (*pgdx*), which increased in the mutant relative to the wildtype starting at T4. By T5, *pgdx* showed a 2.1-fold-change increase in the mutant (p <0.001) (Fig. 4). There was no differential expression (fold change ≥2) observed for other antioxidant genes (*sod*, *prx*) between the strains, however there was a thioredoxin-dependent 2-cys peroxiredoxin, (locus tag BH695_RS02415), which showed fold change increases of ≥1.5 but < 2 (FDR p <0.0001) from T6-T8 in the mutant (Fig. S6).

**FIG 4.**
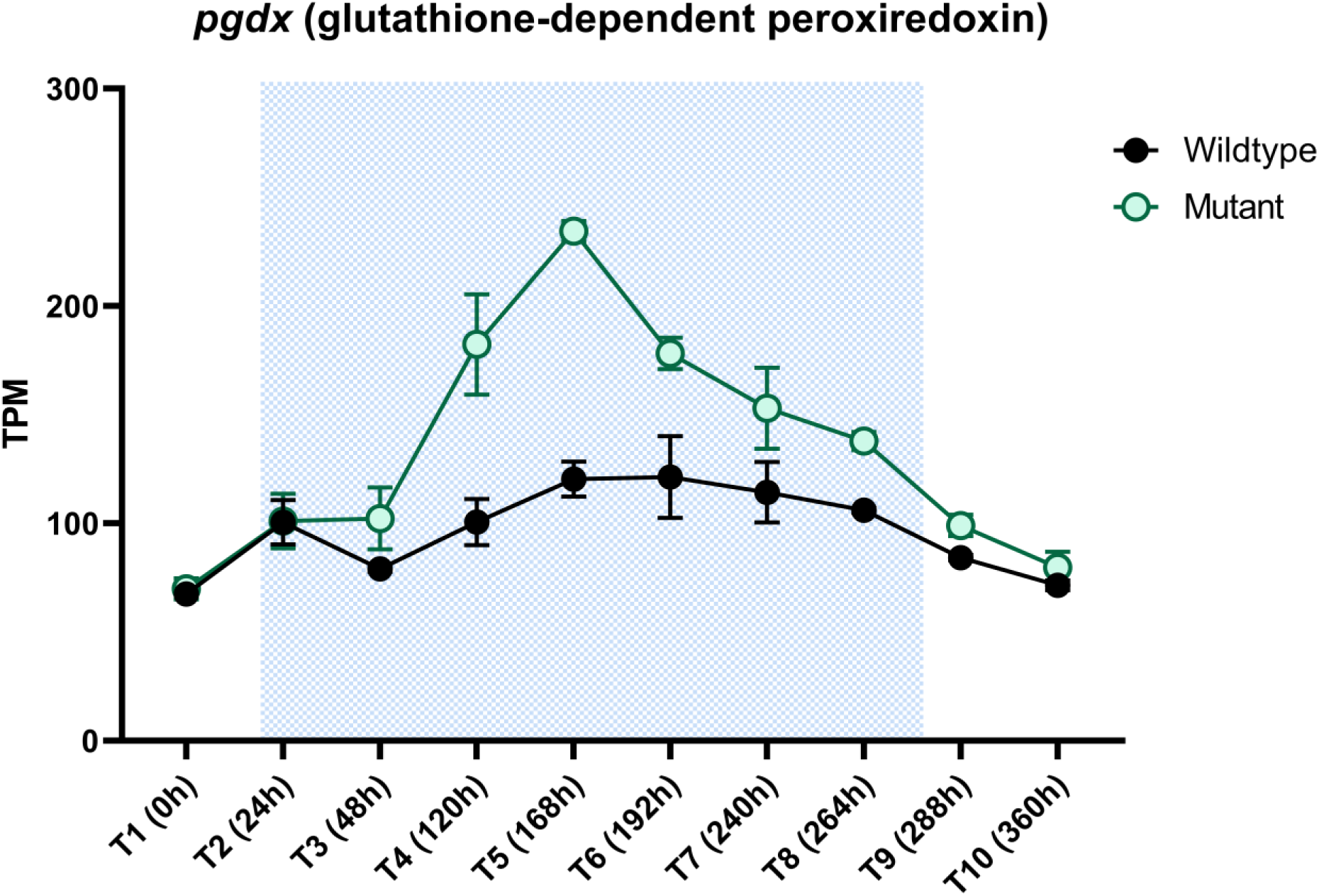
Average TPM data for the mutant (green circles) and wildtype (black circles) for the glutathione-dependent peroxiredoxin, *pgdx*. At T5 the mutant shows has a 2.14 fold change increase (p < 0.0001) in transcript abundance of *pgdx*.

**FIG 5.**
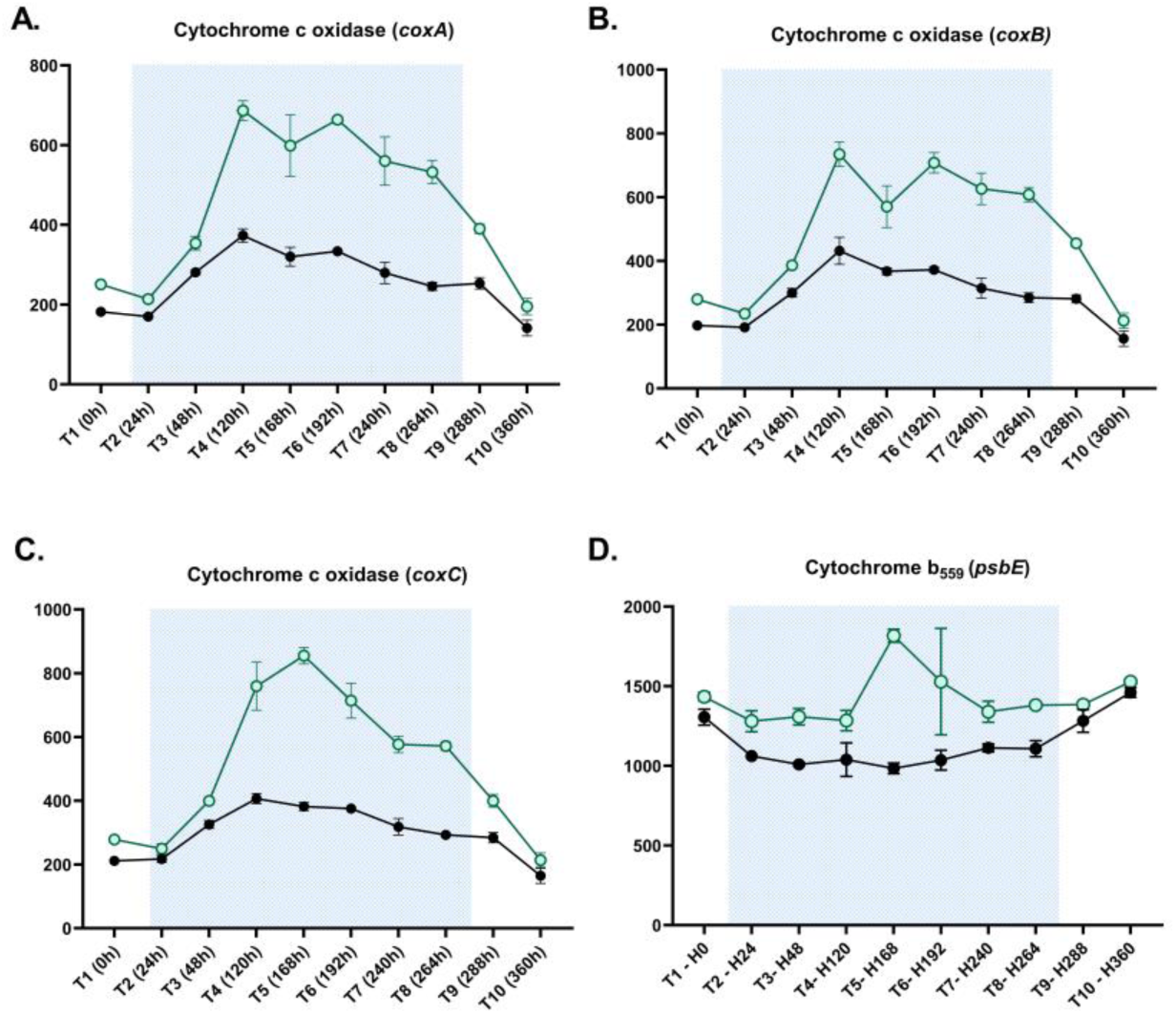
Average TPM data for the mutant (green circles) and wildtype (black circles) for cytochrome c oxidase, *coxBAC*. The x-axis shows each RNA-seq time point in order, from time 0 to 360 hours. (A) Starting at T4, *coxA* shows a fold change increase of 2.1 (p < 0.0001). This fold change increase >2 of *coxA* is seen through the rest of the cold treatment period (T5-T8) (Supplementary tables dataset). (B) Starting at T4, *coxC* shows a fold change increase of 2.1(p<0.001) in the mutant. This fold change increase >2 of *coxC* is seen through the rest of the cold treatment period (T5-T8) (Supplementary tables dataset). (C) At T6, *coxB* shows fold change increase of 2.15 (p < 0.0001) in the mutant, fold change >2 of this gene is maintained through T7 and T8 in the mutant (supplementary tables dataset). (D) In the mutant, cytochrome b_559_ (*psbE*) saw a fold change increase of 2.04, with p < 0.0001 at T5.

In the mutant, a subunit of the photoprotective cytochrome b_559_ complex in PSII (*psbE)* and the respiratory pathway based on cytochrome c oxidase (*coxBAC*) showed differential expression during cold treatment (Fig. 5A-D). From T4 to T8, cytochrome c oxidase expression (*coxBAC*) increased in the mutant (Fig. 5A-C, Tables S6-10), while cytochrome b_559_ alpha subunit (*psbE*), an electron sink, increased ∼2-fold, (p <0.001) at T5 (Fig. 5D). Based on *psbE* and *coxBAC* expression, the mutant had increased expression of thylakoid-membrane-associated electron sinks during cold acclimation.

We also saw changes in expression for genes *pntA* (2 copies), *pntB, petJ*, and *petE*, all of which are associated with proton/electron transporting processes. The genes *pntA* (both copies), *pntB* and *petJ* did not meet our criteria of fold change ≥2, instead they showed fold changes ≥1.5 but <2, with FDR-p values <0.0001 from T6-T8 (Fig. S7, and S8A). The electron transporter, plastocyanin (*petE*), showed fold change decreases >2 (FDR-p < 0.0001) from T7-T9 in the mutant (Fig. S8B).

## Discussion

The purpose of this study was to determine if changes in microcystin production provided an advantage to *Microcystis aeruginosa* PCC 7806 in its response to cold stress. Here we used cold acclimation under constant light – conditions known to generate oxidative stress in phototrophs (33–35). We compared the response of non-toxigenic PCC 7806 Δ*mcyB* (32) with its corresponding toxic wild-type. These strains were chosen to allow comparison within the most genetically similar backgrounds (e.g., 36). We note that over time, genetic changes have accumulated between the two strains (37). As part of this study, we validated that genes individually reported herein as differentially expressed were consistent in terms of genomic structure between strains. Genes for which differential rearrangements or mutations were identified were not considered in our gene expression analysis within the main text or supplemental figures. The physiology of the wildtype and mutant strains were monitored with various methods during growth in chemostats shifted from 26° C to 19° C to induce cold stress (vis a vis 6), a condition known to induce microcystin production. By pairing the temporal RNA-seq analyses with photo-physiology and ROS measurements, we have identified differences in how a toxigenic *vs.* non-toxigenic strain responds and recovers to cool temperature exposure. We present these observations couched within a framework that describes how microcystin influences cell physiology and distinguishes between a defensive role microcystin may play in redox-homeostasis vs. the mutant’s active metabolic restructuring to increase electron sink capacity and degrade ROS.

Microcystin has been implicated as playing a role in the oxidative stress response, due to its thiol-binding abilities (19, 26). Oxidative stress can be generated by photosynthetic processes in cyanobacteria, where disruptions in the redox state of the thylakoid membranes can increase ROS generation if electron source exceeds electron sink capacity. During the cold temperature treatment, we observed a general trend of more reactive thiol groups in the mutant via our CM-H_2_DCFDA assay compared to the wildtype, indicating more ROS damage in that strain (Fig. S3). Interestingly, we also observed an increase in expression of a glutathione-dependent peroxiredoxin (*pgdx*) in the mutant a week (T5) into cold treatment, which coincides with microcystin quota peaking and cell concentration recovery in the wildtype (mutant cold recovery was slightly delayed and started at T6). Glutathione-dependent peroxiredoxins are important for H_2_O_2_ detoxification and have also been implicated as acting as redox sensors (38, 39).

Differential expression of *pdgx* in the mutant during cold treatment is indicative of intracellular stress conditions absent in the wildtype. While other ROS scavengers, such as superoxide dismutase and other peroxiredoxins, behaved similarly in the mutant and wildtype (no fold change >2), the behavior of *pgdx* might indicate more specific regulatory activity than just ROS detoxification alone. Thus, while ROS protection may be one role of microcystin in the cell, the specifics as to the type or location of ROS-inducing damage (H_2_O_2_ v. O^-^) needs more attention.

Glutathione-dependent peroxiredoxins (or glutaredoxins) are reliant on glutathione reductase activity, whose disulfide reductase activity is powered by NADPH (22). Microcystin is capable of binding to glutathione reductase *in vivo*, and it has been hypothesized this binding might either aid or interfere in the oxidative stress response (19, 26). While it is possible the presence of microcystin could be altering the activity of *pgdx* in the wildtype, broader implications of microcystin interference with the glutathione/thioredoxin networks might mean that less NADPH is required for cold metabolism in the wildtype, a scenario consistent with the decreased F_v_/F_m_ beginning at T7. This reduction in F_v_/F_m_ came with no impediment to wildtype growth. On the other hand, the mutant not only showed increased F_v_/F_m_ later into the cold treatment, but there was also an increase in transcription of an NADPH transhydrogenase (*pntAB*) in the mutant relative to the wildtype from T5 through T8 (Fig. S7). This may indicate the mutant takes more action to properly maintain NAD(P)+/NAD(P)H pools during cold growth, which could signify differences in redox coupling of these cofactors and intracellular redox homeostasis in the mutant (40).

We saw additional transcriptional and photo-physiological evidence of metabolic “re-wiring” in the mutant to acclimate to the cold, around the same time *pgdx* gene expression peaked. The mutant upregulated cytochrome b559 (*psbE*) and cytochrome c oxidase (*coxBAC*) in what we interpret as an effort to increase electron sink capacity. Cytochrome b_559_ has been shown to aid in photoprotection, by acting as an alternate electron pathway (41). Cytochrome c oxidase (*coxBAC*) is a respiratory terminal oxidase that generates ATP by consuming oxygen and NAD(P)H reducing equivalents (42). However, cytochrome c oxidase serves a dual role in the cell, both increasing ATP generation and having a photoprotective role, where it acts as an additional electron sink by diverting electrons away from the donor side of photosystem I via plastocyanin or cytochrome c_6_ (42, 43). Because of this, it is thought that activity of this complex in the light may be regulated by the redox state of the electron carriers plastocyanin and cytochrome c_6_ (44). Differential expression of these two electron transporters was observed in the mutant during the cold treatment (Fig. S8A-B). In terms of photoprotective abilities, there was an inverse trend of the *pgdx* expression with cytochrome c oxidase and cytochrome b_559_ expression, giving more credibility to the hypothesis that they are upregulated to help modulate thylakoid redox state in the mutant. Additionally, cytochrome c oxidase relies on NAD(P)H for proton translocation, which further supports an increased need (or may function as a sink) for NAD(P)H in the mutant during cold temperature growth.

Based on transcriptional data presented, we conclude that the mutant and wildtype allocate excitation energy differently. Our NPQ data support this. The mutant had higher NPQ at all incident PAR at T5 compared to the wildtype. This means that during cold growth, the mutant would be more reliant on photo-protective energy dissipation through non-photochemical quenching, which can be achieved either by the orange carotenoid protein in photosystem II or state transitions (45, 46). Increasing electron sinks and induction of NPQ processes in the mutant would decrease generation of ROS, which can result from an over-reduced thylakoid membrane (47, 48). This is interesting, as the excitation pressure of PSII significantly increases in the wildtype at T5 (compared to controls T1 and T10) (Fig. S3). However, the wildtype manages this without increasing alternate electron pathways, or in the short term without increasing NPQ, and still recovers during cold growth despite reduced F_v_/F_m_ after this time. When taken together, these findings show that the mutant and wildtype have different energy dissipation capabilities during cold growth, which may affect NAD(P)+/NAD(P)H pools or the thylakoid membranes.

The absence of microcystin in the mutant increases the need for electron dissipating pathways, indicating microcystin may play a role in processes related to the photosynthetic electron transport chain. Complementary to this idea, if the mutant employs these strategies to mitigate or reduce ROS production, the absence of “turning on” these strategies in the wildtype could mean microcystin is in fact playing a role in the oxidative stress response, as has been previously hypothesized by other researchers.

### Microcystin production alters cold stress management strategies

In response to cold treatment (11 days), the non-toxigenic mutant upregulated a glutathione-dependent peroxiredoxin and a cytochrome c oxidase complex, both of which help mitigate or reduce ROS generation in cyanobacteria. These responses temporally coincided with an increase of microcystin in the wildtype. Coincident in time, photo-acclimation strategies differed, in which the strains showed opposite response in F_v_/F_m_, and NPQ measurements increased in the mutant. This implies cold acclimation alters electron transport capabilities differently in the mutant and wildtype strains. Our NPQ measurements also indicate inherent differences in the redox state of the thylakoid membranes between the strains during cool temperatures (14).

Indeed, differences in redox-related proteins have been shown to differ between these strains, and conditions that can cause over reduction of the thylakoid membranes, such as high-light growth, provide the toxigenic strain an advantage (18, 49, 50). Collectively, our observations lead to the idea that increased microcystin per-cell during the cold treatment may provide a longer-term benefit (days versus hours) during conditions which can stress photosynthetic metabolism resulting in more ROS generation (here brought on by cold temperature growth). This leads us to our hypothesis that in locations that experience seasonal temperature trends like Lake Erie, microcystin quota may be higher under conditions where solar days are longer, and the water is clearer and colder (*e.g.,* June). This phenomenon may be overlooked by total microcystin measurements (μg/L) alone, as overall biomass is lower earlier in the growing season. The replacement of microcystin producers by non-producers in August (where temperatures are warmer, solar days shorter and water less clear) occurs during periods that may favor photo acclimation strategies that differ from those seen during high light and low temperature growth periods. This would include the use of enzymes that directly degrade reactive oxygen (*e.g.,* like peroxiredoxin), noting that specific strategies may be tailored to certain reactive oxygen species. Thus, there could be a connection between photo acclimation strategies for *Microcystis*, the nature of the photosynthetic stressors, and succession of toxigenic genotypes of *Microcystis*. We hypothesize that trade-offs in the differing strategies of managing photosynthetic stressors may therefore play an important role in regulating/selecting the toxigenic composition of *Microcystis* blooms (although other factors are also likely playing a role) (51). Agent-based modeling has suggested that nitrogen availability may contribute to the succession of toxigenic to non-toxigenic strains in Lake Erie (52). Data from our experiment can be integrated into models such as this, allowing us to quantify how nutrients and temperature contribute to genotypic succession. Doing so may help improve management and mitigation strategies of *Microcystis*-dominated harmful algal blooms. Yet *Microcystis* blooms are complicated: the influence(s) of other environmental factors as well as competition with biology needs to be appropriately accounted for in our stewardship of this systems.

## Methods

### Continuous cultures

*M. aeruginosa* PCC 7806 was obtained from the Pasteur Culture Collection and has been maintained for the last decade as both active cultures and cryopreserved stocks. *Microcystis aeruginosa* PCC7806 *ΔmcyB* was generated by Dittmann and colleagues (32) and kindly provided by Professor E. Dittmann in 2016 (University of Potsdam, Germany). To assess gene mutations or transposon insertion/disruption (53), we re-sequenced and closed the genomes of both strains. Any differentially expressed genes discussed in this paper have been examined (along with adjacent intergenic spacer regions) to ensure phenotypic differences were not due to gene mutations occurring during this experiment (37). We refer to these isolates as “wildtype” and “mutant”. Chemostat design and construction followed (6). Duplicate 1-L chemostats were established for each strain using modified CT medium (54), with nitrogen (N) concentrations at 323 μM supplied as a mix of Ca(NO_3_)_2_ and KNO_3_ (N molar ratio of 1.37:1) (6). Phosphorus (P) was supplied as 16.3 μM by K_2_HPO_4_, resulting in N:P molar ratios of ∼20. Chemostats were maintained in the same incubator at a measured light intensity of ∼50 µmol photon m^-2^ s^-1^, with a dilution rate of 0.25 d^-1^ controlled by independent peristaltic pumps. Chemostats were acclimated to 26° C, (referred to as the “warm” treatment), and 50 µmol photon m^-2^ s^-1^ for ∼3 weeks, to ensure steady-state was achieved. After chemostat cell concentrations remained constant (± 8%) for about a week, incubator temperature was lowered to 19° C, (referred to as the “cold” treatment) and maintained at 19° C for 11 days before it was returned to 26° C. Re-acclimation to 26° C (referred to herein as “recovery”) was monitored for one week. Sample collection (details below) occurred around 12:00 PM daily. Light was kept constant throughout the experiment. Temperature was monitored every 15 min with a Hobo Tidbit TempLogger (Onset Computer Corporation) to account for potential temperature fluctuations. At the end of the experiment, chemostat cultures were assessed for contamination by transferring aliquots to LB medium (55) followed by incubation at 26° C in the dark.

### Cell abundance and size

Culture (1 ml) was collected from each chemostat through a sampling port using a sterile syringe. The samples were diluted ten-fold in fresh CT medium and assayed with a Guava EasyCyteHT flow cytometer (Millipore). Cell concentrations were determined by gating on red fluorescence, a proxy for chlorophyll *a*, and forward scatter, a proxy for size proxy (6).

### Measurement of microcystin

Microcystin concentrations were estimated from 50 mL of culture collected daily. Samples were filtered onto 47-mm glass fiber filters (GF-75, Advantec) *via* vacuum filtration, immediately flash-frozen in liquid nitrogen, then stored at -80° C until processing. Quantification of microcystin and congeners was completed by LC-MS as outlined in detail at *protocols.io* (56).

### Measurements of photosynthetic physiology

To assess changes in photo-physiology, 3-mL of chemostat culture was collected daily. Rapid light curves (RLC) using a Pulse-Amplitude Fluorometry (Phyto-PAM II, Heinz Walz GmbH) were generated after 60 s of low-light far-red adaptation. Saturation pulse kinetics were checked to ensure a fluorescence peak was achieved before sample processing. For rapid light curves, saturating pulses (5,000 µmol photon m^-1^ s^-2^) were applied every 30 s after actinic light exposure. The RLC consisted of 12 actinic light steps, starting at a PAR (µmol photon m^-1^ s^-2^) of 1 and ending at 1275 PAR (µmol photon m^-1^ s^-2^). From these rapid light curves, maximum potential quantum efficiency (F_v_/F_m_), nonphotochemical chlorophyll fluorescence quenching (NPQ), and photosystem II excitation pressure (1-qP) estimates were calculated by the Phyto-PAM II software.

### Oxidative stress assessment

A working concentration of approximately 1.15 x 10^7^ cells were used for the oxidative stress assays (CM-H_2_DCFDA). Cells were first pulled from the chemostats and pelleted by centrifugation at 3,700 RCF for 15 min at 19° C. Cell pellets were washed and re-suspended in PBS buffer free of KCl and incubated in the dark with 25-μM of 2’,7’-dichlorodihydrofluorescein diacetate (CM-H_2_DCFDA) suspended in dimethylsulfoxide (DMSO) for 60 min at room temperature (Molecular Probes, Invitrogen). After incubation, cells were washed again and resuspended in PBS buffer. Changes in green fluorescence were measured *via* flow cytometry using a Guava easyCyteHT flow cytometer (ex_λ_ = 485 nm, em_λ_ = 535 nm). Each sample was run for 120 seconds, or until a maximum of 5000 cells were counted. Control culture green fluorescence (cells without CM-H_2_DCFDA) was subtracted from cultures incubated with CM-H_2_DCFDA, for net fluorescent gains. Green fluorescence emission is indicative of the thiol-reactive chloromethyl group of the CM-H_2_DCFDA probe reacting with glutathione and other thiols (Invitrogen).

### RNA extraction

Samples (25 mL) were collected daily on 47-mm diameter, 0.2 µm nominal pore-size polycarbonate filters and immediately flash-frozen in liquid nitrogen before storage at -80° C. We selected ten points across the time course (four chemostats x 10 time points, n=40) for RNA sequencing based on cell concentration observations (Supplementary Table 1). To extract RNA, we used the acid-phenol chloroform bead-beating method described in (57) and Turbo DNA-free kit to eliminate contaminating DNA (Invitrogen). Checks for contaminating DNA were completed by PCR (using 515F and 806R primers targeting the 16s rRNA gene) and visualization on an agarose gel. Where DNA contamination was present (*i.e.,* PCR products observed in the gel), the sample was processed with another round of DNAse and rechecked until no band was present. Final RNA concentrations were determined using a high-sensitivity Qubit RNA assay per manufacturer’s instructions (Invitrogen).

### RNA sequencing and sequence analyses

RNA samples were sent to Discovery Life Sciences (previously Hudson Alpha, Huntsville, AL) where sample library prep and Illumina Novaseq 6000 platform sequencing (50 million reads, 150 bp, paired end) were performed after ribosomal RNA reduction. A TruSeq rRNA reduction was performed on all samples *ex-silico*, according to manufacturer protocol. Sequence data was loaded to CLC genomics Workbench (v. 21.0.4) where it was interleaved, filtered, and trimmed using default program settings. Due to the axenic nature of the chemostats, transcriptome RNA-sequencing reads were mapped (length fraction 0.8, similarity fraction 0.9) to the rRNA genes (PCC7806 has two sets of 5S, 16S and 23S genes) from the *Microcystis aeruginosa* PCC7806SL genome to remove ribosomal reads (58). Reads that did not map to rRNA genes were then stringently mapped (length fraction 0.9, similarity fraction 0.9) to the annotated *Microcystis aeruginosa* PCC 7806SL genome (GenBank accession NZ_CP020771.1, annotated Jan-28-2020) (58), and normalized to transcripts per million (TPM). Differential expression analysis within each time point (calculated as expression of wildtype vs. expression of mutant at each time point) was performed using CLC’s differential expression algorithm. Genes were considered differentially expressed if FDR-corrected p-values were ≤ 0.05 and fold change was ≥ 2. In our supplemental materials, we considered genes with fold changes ≥ 1.5 and FDR-p values < 0.05 for further investigation into genes that had fold changes ≥ 2 and FDR p-values < 0.05 in the main text. For downstream analysis of gene function, coding sequences (CDS) from *Microcystis aeruginosa* PCC7806SL (NZ_CP020771.1, annotated Jan-28-2020) were supplemented/merged with annotations from KEGG BlastKOALA. Genes of interest (displayed fold change ≥ 2) that were annotated as “hypothetical genes” were run through NCBI’s conserved domain protein search and HMMER for protein homology. Reads mapping to genes that were not annotated and that failed to match with a homologous protein search were omitted from downstream analyses. Tables listing differentially expressed genes by RNA-seq time points (T1-T10) are provided in supplemental tables S3-S12. For this study, we discuss results in terms of transcript abundance: we define that here as “relative abundance” within the normalized transcriptomes from each time point.

### Data visualization and statistical analyses

All statistical analyses were performed using R studio (4.2.1), PRIMER (v.7), or GraphPad Prism (v. 8.0.2). To check sphericity of the data for repeated measures ANOVA Phyto-PAM measurements were analyzed using the Mauchly test in R. If the data did not fit assumptions (p-value from Mauchly test was < 0.05), the Geisser Greenhouse correction was used (59).

Repeated measures and two-way ANOVA were run in GraphPad Prism (v. 8.0.2). GraphPad Prism was used for time point comparisons of the physiological measurements (cell counts, F_v_/F_m_ and NPQ) within or between the strains. PRIMER (v.7) was used for non-metric multidimensional scaling (nMDS). Square-root-transformed TPM expression of all genes in the PCC7806SL genome (4,889) was used in calculating the similarity matrix used in the nMDS.

## Data availability

All raw paired-read data used in this study is publicly available at NCBI under the BioProjectID PRJNA1008692.

## Acknowledgements

This work was supported by funds from the National Science Foundation (OCE-1840715) and the National Institute of Environmental Health Sciences (1P01ES028939–01) through funds provided to the Great Lakes Center for Fresh Waters and Human Health at Bowling Green State University, and the National Oceanographic and Atmospheric Administration (NA18NOS4780175). We also thank the generous support of the Kenneth & Blaire Mossman Endowment to the University of Tennessee.

## Contributions

Conceptualization: G.F.S., R.M.M, S.W.W.; Data curation: G.F.S.; Formal Analysis: G.F.S., S.W.W.; Funding acquisition: S.W.W., F.L.W, G.S.B.; Investigation: G.F.S., L.E.S, B.W.; Methodology: G.F.S., R.M.M., S.W.W.; Project administration: S.W.W.; Resources: S.W.W., G.L.B.; Supervision: S.W.W., G.L.B.; Validation: G.F.S.; Visualization: G.F.S.; Writing – original draft: G.F.S.; Writing – review & editing: all authors.

The authors declare no conflict of interest.

